# Prevalence, complete genome and metabolic potentials of a phylogenetically novel cyanobacterial symbiont in the coral-killing sponge, *Terpios hoshinota*

**DOI:** 10.1101/2021.02.04.429686

**Authors:** Yu-Hsiang Chen, Hsing-Ju Chen, Cheng-Yu Yang, Jia-Ho Shiu, Daphne Z. Hoh, Pei-Wen Chiang, Wenhua Savanna Chow, Chaolun Allen Chen, Tin-Han Shih, Szu-Hsien Lin, Chi-Ming Yang, James Davis Reimer, Euichi Hirose, Budhi Hascaryo Iskandar, Hui Huang, Peter J. Schupp, Chun Hong James Tan, Hideyuki Yamashiro, Ming-Hui Liao, Sen-Lin Tang

**Author notes:** Yu-Hsiang Chen and Hsing-Ju Chen contributed equally to this work. Author order was determined on the basis of contribution. Corresponding author Corresponding author information: Dr. Sen-Lin Tang, Biodiversity Research Center, Academia Sinica, Address: A305, Interdisciplinary Research Building for Science and Technology, 128 Academia Road, Sec. 2, Nankang, Taipei 11529, Taiwan.

## Abstract

*Terpios hoshinota* is a ferocious, space-competing sponge that kills a variety of stony corals by overgrowth. Outbreaks of this species have led to intense coral reef damage and declines in living corals on the square kilometer scale in many geographical locations. Our large-scale 16S rRNA gene survey across three oceans revealed that the core microbiome of *T*. *hoshinota* included operational taxonomic units (OTUs) related to *Prochloron*, *Endozoicomonas*, *Pseudospirillum*, SAR116, *Magnetospira*, and *Ruegeria*. A *Prochloron*- related OTU was the most dominant cyanobacterium in *T*. *hoshinota* in the western Pacific Ocean, South China Sea, and Indian Ocean. The complete metagenome-assembled genome of the *Prochloron*-related cyanobacterium and our pigment analysis revealed that this bacterium had phycobiliproteins and phycobilins and lacked chlorophyll *b*, inconsistent with the iconic definition of *Prochloron*. Furthermore, the phylogenetic analyses based on 16S rRNA genes and 120 single-copy genes demonstrated that the bacterium was phylogenetically distinct to *Prochloron*, strongly suggesting that it should be a sister taxon to *Prochloron*; we therefore proposed this symbiotic cyanobacterium as a novel species under a new genus: *Candidatus* Paraprochloron terpiosii. With the recovery of the complete genome, we characterized the metabolic potentials of the novel cyanobacterium in carbon and nitrogen cycling and proposed a model for the interaction between *Ca.* Pp. terpiosi LD05 and *T. hoshinota.* In addition, comparative genomics analysis revealed that *Ca.* Paraprochloron and *Prochloron* showed distinct features in transporter systems and DNA replication.

**Importance:** The finding that one species predominates cyanobacteria in *T*. *hoshinota* from different geographic locations indicates that this sponge and *Ca*. Pp. terpiosi LD05 share a tight relationship. This study builds the foundation for *T*. *hoshinota*’s microbiome and paves a way for understanding the ecosystem, invasion mechanism, and causes of outbreak of this coral-killing sponge. Also, the first *Prochloron*-related complete genome enables us to study this bacterium with molecular approaches in the future and broadens our knowledge of the evolution of symbiotic cyanobacteria.

## Introduction

The coral-killing sponge, *Terpios hoshinota*, has received attention since its outbreaks were detected in several coral reefs in the western Pacific Ocean, South China Sea, and Indian Ocean (1–7). The encrusting sponge can kill various stony corals by overgrowth (8). It grows up to 23 mm per month (9), and its fast growing and competitive nature enables it to kill scleractinian corals quickly and at a high rate (30–80%) across biogeographic regions (4, 5, 7, 10). For instance, *T*. *hoshinota* overgrowth killed 30% of corals in some coral reefs of Green Island, Taiwan in just one year (2). Similarly, the sponge jeopardizes coral reefs in numerous regions of Indonesia, Malaysia, Japan, India, and the Great Barrier Reef (3, 10–12). The gradual spread of *T. hoshinota* makes it a serious threat to coral reefs. However, little is known about what causes its outbreaks.

Sponges are commonly known to harbor complex microbial communities, and symbiotic microorganisms play vital roles in the development, health, and nutrient acquisition of their hosts (13). The microbe and host form an ecological unit called a holobiont. In *T. hoshinota*, the sponge associates with a bacterial community of relatively low diversity that is dominated by cyanobacteria (14). Ultrastructural observations have clearly shown that cyanobacteria are densely distributed in the sponge, contributing to 50% of the total cell volume (1, 14, 15). The blackish color of *T. hoshinota* is also attributed to cyanobacteria (14). Accordingly, the sponge is called cyanobacteriosponge. Several studies have shown that cyanobacteria play important roles in the sponge’s growth and competition with corals. For instance, high numbers of cyanobacteria can be observed in the sponge’s larvae (16, 17), suggesting that they are transmitted vertically during embryogenesis in this particular sponge (18). Second, *in situ* light shading was shown to discontinue *T. hoshinota* expansion (19). Even short-term shading can cause a long-term decrease in the biomass of symbiotic cyanobacteria and lead to irreversible damage to *T. hoshinota* (20). Furthermore, one study found that, when *T. hoshinota* encountered certain corals, the sponge formed a hairy tip structure packed with dense cyanobacteria to interact with the corals (8). These results indicate that cyanobacteria are vital for its overgrowth of corals.

Though important, the identity of the symbiotic cyanobacteria is still unclear. In 2015, Yu *et al*. isolated and cultivated *Myxosarcina* sp. GI1, a baeocytous cyanobacterium, from *T. hoshinota* at Green Island (21). However, electron microscopy has not identified vegetative cell aggregates of baeocyte, a type of reproductive cell, in *T. hoshinota* (14–16). Moreover, our previous study analyzed 16S ribosomal RNA gene sequences from Green Island and found that the dominant cyanobacterium in *T. hoshinota* is a novel species closely related to *Prochloron* sp. (14). *Prochloron*, a genus comprises of a single species, is an obligate symbiont of some ascidians; the hallmarks and definitions of this genus include the present of chlorophyll *b* and the lack of phycobilins, which is unusual in cyanobacteria (15). However, a pigment analysis reported that the cyanobacteria in *T. hoshinota* contained phycobilins. Hence, the identity and characteristics of the predominant cyanobacterium in *T. hoshinota* remain uncertain, as does the question of whether the predominance of this particular cyanobacterium is present in all the Indo-Pacific regions. On the other hand, since cyanobacteria are attributed to the sponge’s health and invasion capacity, the ecological relationship and molecular interaction between *T. hoshinota* and its cyanobacteria needs to be determined.

Other bacteria in the *T. hoshinota* microbiome also contribute to holobiont functions. Microbial community in sponge can be shaped by host-related factors, such as the immune system and nutrient exchanges (22, 23). Strict host selective pressure can stabilize the microbial community (24). On the other hand, the community can also be determined by environmental factors, such as light availability, pH, and temperature (22). Nevertheless, no study has investigated the microbiome of *T. hoshinota* under different biogeographical backgrounds. Hence, this study investigated the bacterial community *T. hoshinota* across different oceans to examine whether the bacterial community is stable or depends on geography.

Most sponge symbionts, including the dominant cyanobacteria in *T. hoshinota*, are uncultivable. Metagenomic methods can be used to determine the microbial composition in *T. hoshinota*, and knowing this composition will help us infer the ecosystem of the *T. hoshinota* holobiont. In this study, we conducted 16S rRNA gene survey to investigate the bacterial composition and diversity of *T. hoshinota* samples from a wide geographical range across the western Pacific Ocean, South China Sea, and Indian Ocean. Knowing the predominance of a cyanobacterium was ubiquitous, the complete genome of this cyanobacterium was reconstructed by whole-genome shotgun Nanopore and Illumina sequencing. The following genomic and comparative genomic analyses elucidated the phylogenetic affiliation and taxonomy of the dominant symbiotic bacterium in *T. hoshinota* and putative symbiotic interactions between the bacterium and the host.

## Results

### Bacterial diversity in *Terpios hoshinota* samples from different oceans

To characterize the bacterial communities in coral-encrusting *T. hoshinota* across different geographical backgrounds, samples were collected from 15 locations in the Pacific Ocean, South China Sea, and Indian Ocean (Table S1). 16S ribosomal RNA (rRNA) gene sequencing was used to determine the bacterial communities.

Statistical tests showed that operational taxonomic unit (OTU) Chao1 richness and Faith’s phylogenetic diversity (PD) indices were not significantly associated with the oceanic regions from which the samples were collected (Chao1: *p* = 0.23, PD: *p* = 0.30) (Fig. S1a and S1b). In terms of Shannon diversity, the indices were also not significantly associated with the oceanic regions when the outliners were removed (Fig. S1c). Some samples—e.g., TWKT, MYRD, and MVFF samples—from the same locations showed high variation in Chao1 and Shannon indices. In contrast to the alpha diversity indices, the similarities in microbial composition were associated with sample origins (Fig. S2). PERMANOVA showed that samples from the same ocean were significantly more similar than those from different oceans (Bray Curtis: pseudo-*F* = 5.13514, *p* = 0.001, Unweighted UniFrac: pseudo-*F* = 1.60401, *p* = 0.01). When relative abundances were taken into consideration, the effect of the sea on similarities increased (weighted UniFrac: pseudo-*F* = 6.09022, *p* = 0.001).

### *Prochloron*-related bacterium dominated *T*. *hoshinota* bacterial communities across different oceans

Totally we recovered 1,164 OTUs. Microbial composition analysis revealed that OTUs related to *Cyanobacteria* and *Proteobacteria* accounted for most of the bacterial composition across the samples, ranging from 81.5–97.4% (Fig. 1 and Fig. S3a). *Bacteroidetes* and *Spirochaetae* were present across all the samples, though their abundances were only 0.34–6.29%. Genus-level analysis showed that a *Prochloron-*related OTU, referred as Prochl-OTU01, was dominant and present in all the samples (Fig. S3b and Table S2). Moreover, the *Prochloron-*related OTU accounted for most of the *Cyanobacteria* in the samples (Table S2).

**Figure 1.**
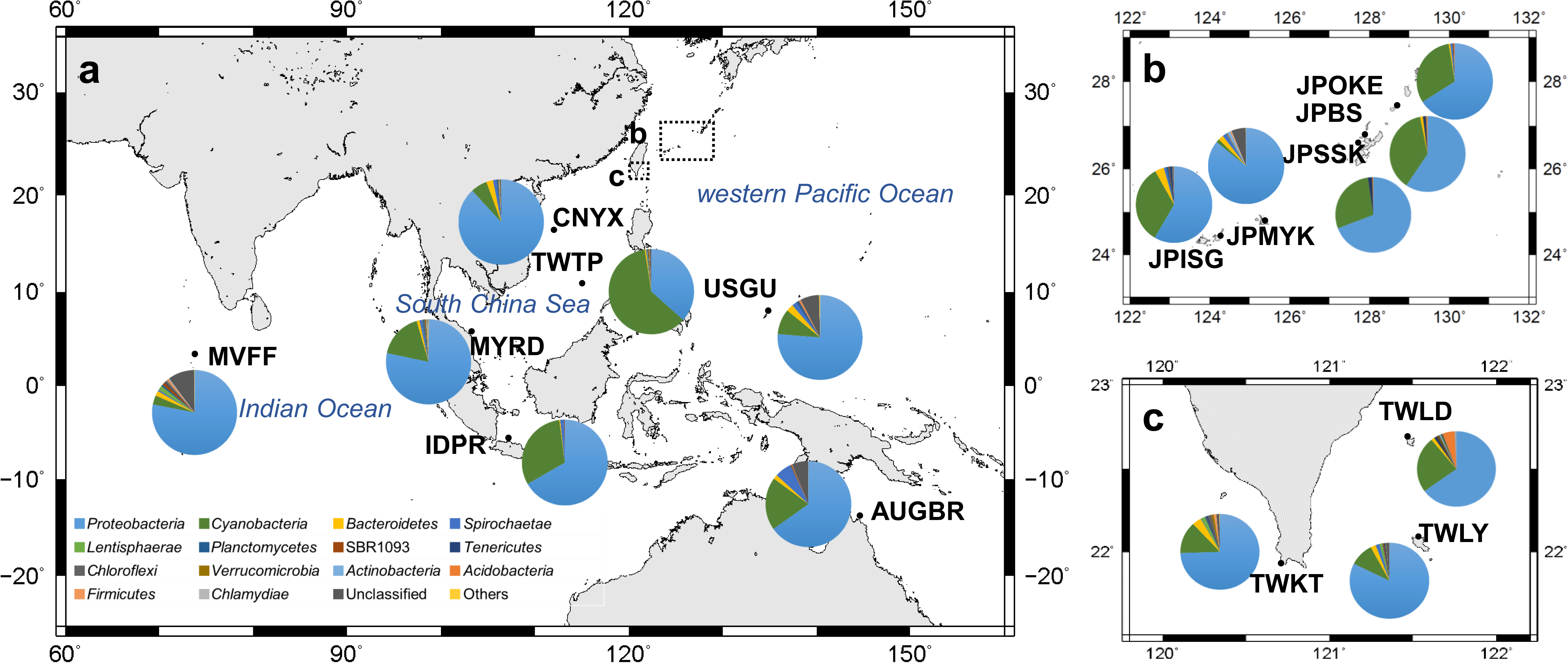
Proportions of *T*. *hoshinota*-associated bacterial communities across oceans. *Proteobacteria* (blue) and *Cyanobacteria* (green) were the dominant groups, ranging from 81.5–97.4% of community. (a) Bacterial compositions across the western Pacific Ocean, South China Sea, and Indian Ocean. Maps on the right show results in Okinawa, Japan (b), and Taiwan (c). Different colors represent different bacterial phyla. Percentage composition of phyla less than 0.05% are grouped as “Others.”

Besides Prochl-OTU01, we also identified other core microbiome members, defined as OTUs that were present at least 90% of the samples. The OTUs of the core microbiome were related to *Endozoicomonas*, *Pseudospirillum*, SAR116, *Magnetospira*, and *Ruegeria.* Consistent with genus-level analysis, these genera were co-occurred across all the samples. This core microbiome averagely contributed 80% of bacterial abundance in each sample (Fig. S3b).

### Structure of symbiotic *Prochloron*-related bacteria in *T*. *hoshinota*

To reveal the detailed structure of the *Prochloron*-related bacteria, a *T*. *hoshinota*- encrusting abiotic substrate was observed using transmission electron microscopy and scanning electron microscopy (Fig. 2). The *T. hoshinota* sponge, roughly 500 μm thick, were supported by the bundle of tylostyle spicules, and the spicules on the surface were interlaced (Fig. 2a-b). Inside the body, spherical bacterial cells 4–6 μm in diameter were widely distributed, and cells were observed in various stages of cell division (Fig. 2c and 2f). A study published in 2015 isolated a baeocytous cyanobacterium, *Myxosarcina* sp., from *T. hoshinota* (21). However, no vegetative cell aggregate of baeocytes could be observed in our results.

**Figure 2.**
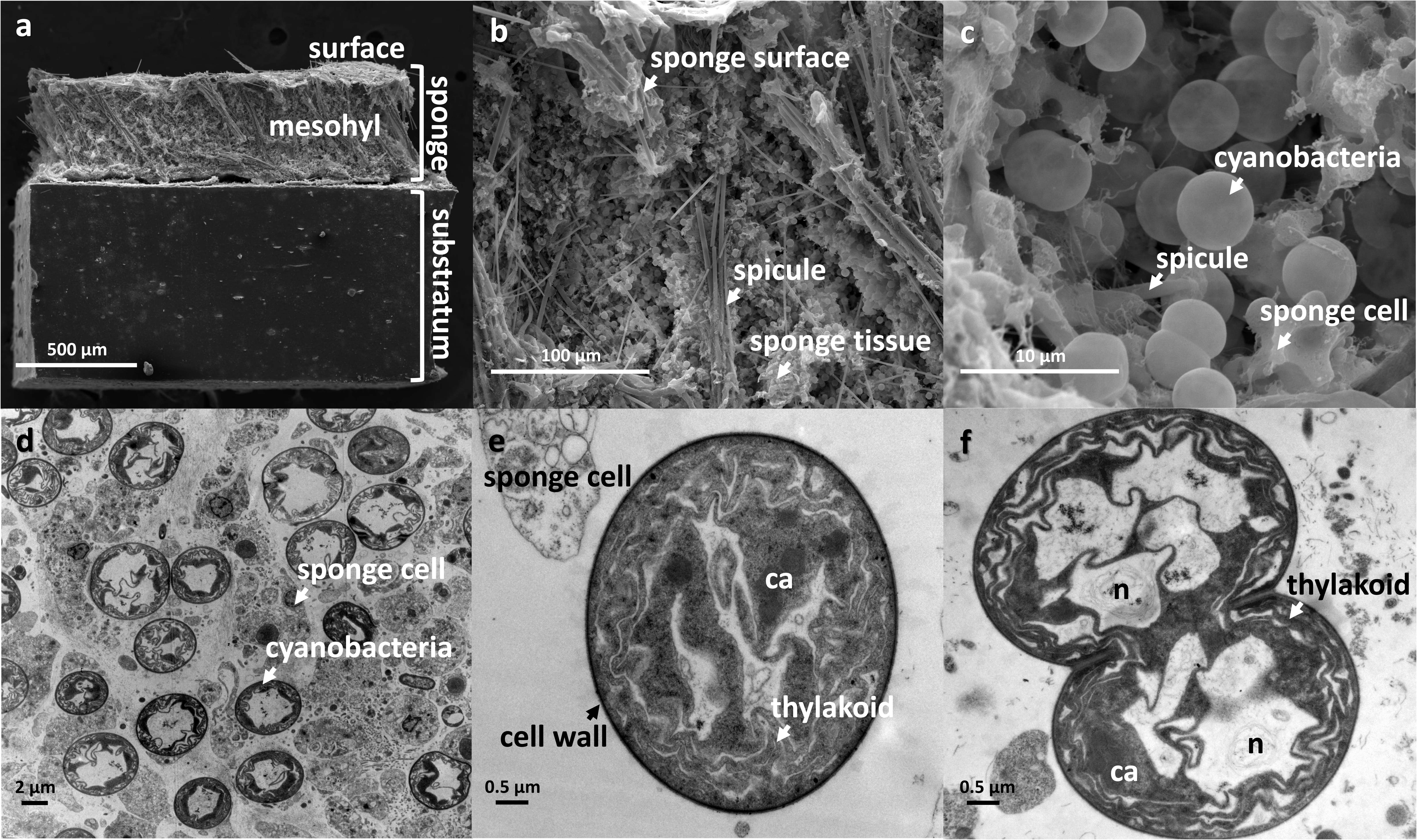
Electron micrographs of *T*. *hoshinota* and associated cyanobacteria. (a-c, SEM; d-f, TEM). (a) Sponge covering an abiotic substratum. The sponge’s skeleton was arranged in a tangential style, and the mesohyl was supported by a tract of spicules. (b, c) Close view of the mesohyl. Numerous spheres composing the inside of the mesohyl, and cross-matched with (d); their thylakoids identified the spheres as cyanobacteria. The detailed structures of the cyanobacteria were observed from a single cyanobacterial cell (e) and a dividing cell (f); they included the cell wall, thylakoid, carboxysome (ca), and nucleoid (n).

TEM results showed that these spherical cells contained thylakoids, typical compartments inside cyanobacteria, and the arrangements were parietal (Fig. 2d–f). In addition, carboxysomes were found; gas vesicles were absent in these cells, which is consistent with a previous description of *Prochloron*’s structure (25).

### Genome assembly of a novel *Prochloron-*related species

The predominance of a *Prochloron*-related OTU in *T*. *hoshinota* from various locations indicates the intimate association between bacterium and host. To reveal the identity and characteristics of the *Prochloron*-related bacterium, a *Prochloron*-related genome, referred as to LD05, was reconstructed by *de-novo* assembly of the metagenome from the sample at Green Island, Taiwan (Fig. 3). We used Nanopore sequencing and metaFlye, a cutting-edge long read metagenome assembler (26), to assemble metagenome and successfully recovered a single circular contig, which was annotated as cyanobacterium without any gap. The complete metagenome-assembled genome (MAG) was then polished with Illumina reads to obtain complete MAG with high accuracy. The mapping coverage of Illumina reads was 99.98% of the MAG and the mean depth was 983 times.

**Figure 3.**
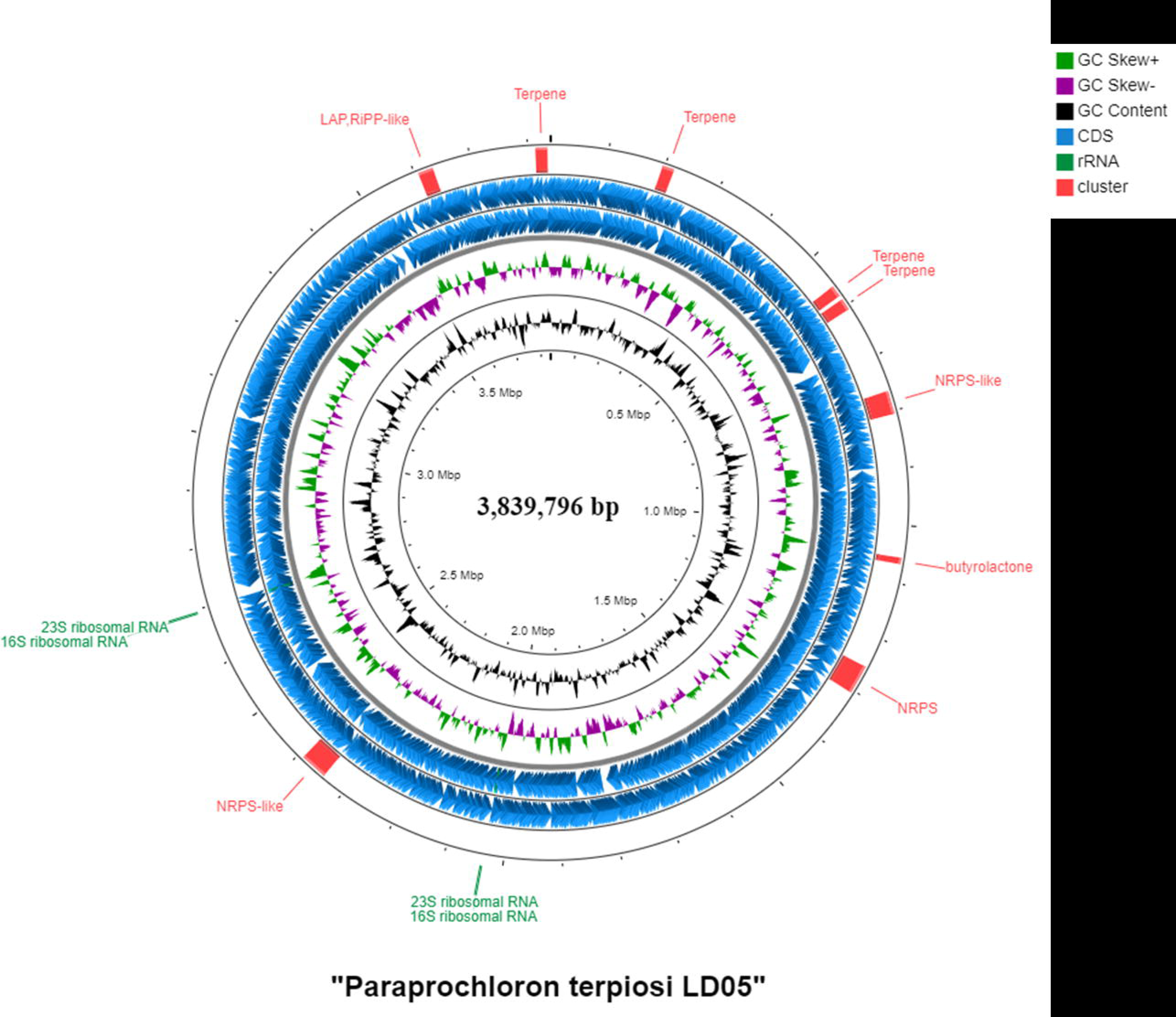
Representation of the “Paraprochloron terpiosi LD05” genome. The genome is 3,839,796 bp. The rings from inside to outside represent GC content (black), GC skew- (purple), GC skew + (green), coding sequence regions (blue), rRNA gene sequences (dark green), and secondary metabolite gene clusters (red).

The complete MAG contained two copies of 16S and 23S rRNA gene sequences (Fig. 3 and Table 1) and 46 tRNA genes; one of the 16S rRNA gene sequence was 100% identical to that of the predominant Prochl-OTU01 identified from the community survey. The analysis of the GTDB-Tk demonstrated that the LD05 MAG was closest to *Prochloron didemni,* but only with 77.97% average nucleotide identity (ANI).

Recently, a putative *Prochloron* species genome called SP5CPC1 was recovered from the metagenome of a sponge microbiome (27). The ANI analysis found that the LD05 genome shared 93.18% identity with the SP5CPC1 genome, which is below the 95% ANI cutoff for species delineation (28). The LD05 shared a similar genome size, GC ratio, and coding density with SP5CPC1 (Table S3). On the other hand, *P*. *didemni* P2-P4 had a much larger genome and lower GC ratio and coding densities (Table S3). There was also a discrepancy between LD05 and *P*. *didemni* P2-P4 based on the average amino acid identity (AAI) and percentage of conserved proteins (POCP) analyses (Fig. S4). The LD05 shared 91% AAI with SP5CPC1, but only 70% AAI with *P*. *didemni* P2–P4 genomes. On the other hand, the POCP between LD05 and *Prochloron* genomes ranged from 49.8–52%, close to a proposed 50% POCP cutoff for the genera delineation (29). Taken together, the LD05 and SP5CPC1 genomes were different species, and both were more distantly related to *P*. *didemni*.

### Phylogenetic and functional genomic analyses revealed distinct characteristics of **LD05 and SP5CPC1 genomes compared to other *Prochloron* genomes**

Although the LD05 and SP5CPC1 genomes had certain degrees of similarity to *Prochloron* genomes, the phylogenetic, genomic, and functional genomic analyses identified distinct characteristics between them and *Prochloron*, suggesting that they belong to a different genus.

A phylogenetic tree of 120 cyanobacteria was constructed based on 120 single-copy genes to determine phylogenic affiliation (Fig. 4a and Fig. S5). The tree demonstrated that the LD05 and SP5CPC1 formed a clade that was sister to the clade containing *P*. *didemni*. The larger clade encompassing the two clades was adjacent to the clade containing *Synechocystis*, *Myxosarcina*, and other cyanobacteria. To provide an in-depth view, homologs of the LD05 16S rRNA gene with high similarities were retrieved from the NCBI database. A tree constructed by these 16S rRNA sequences depicted two distinct clades, referred as to Clade I and Clade II (Fig. 4b). Clade II contained 16S rRNA gene sequences from three genomes of *P*. *didemni* strains and other 16S rRNA sequences from various ascidians. In contrast, Clade I mainly comprised sequences from sponge holobionts, including SP5CPC1, LD05, and 16S rRNA sequences from a variety of other sponges. Although many sequences in Clade I were assigned as *Synechocystis* and *Prochloron* 16S rRNA sequences, we found that they actually shared higher identities with the members of Clade I than *Synechocystis* or *Prochloron didemni*. Moreover, the genome phylogenomic analysis also revealed that the SP5CPC1 and LD05 are phylogenetically distant from cultivated *Synechocystis* (Fig. 4a).

**Figure 4.**
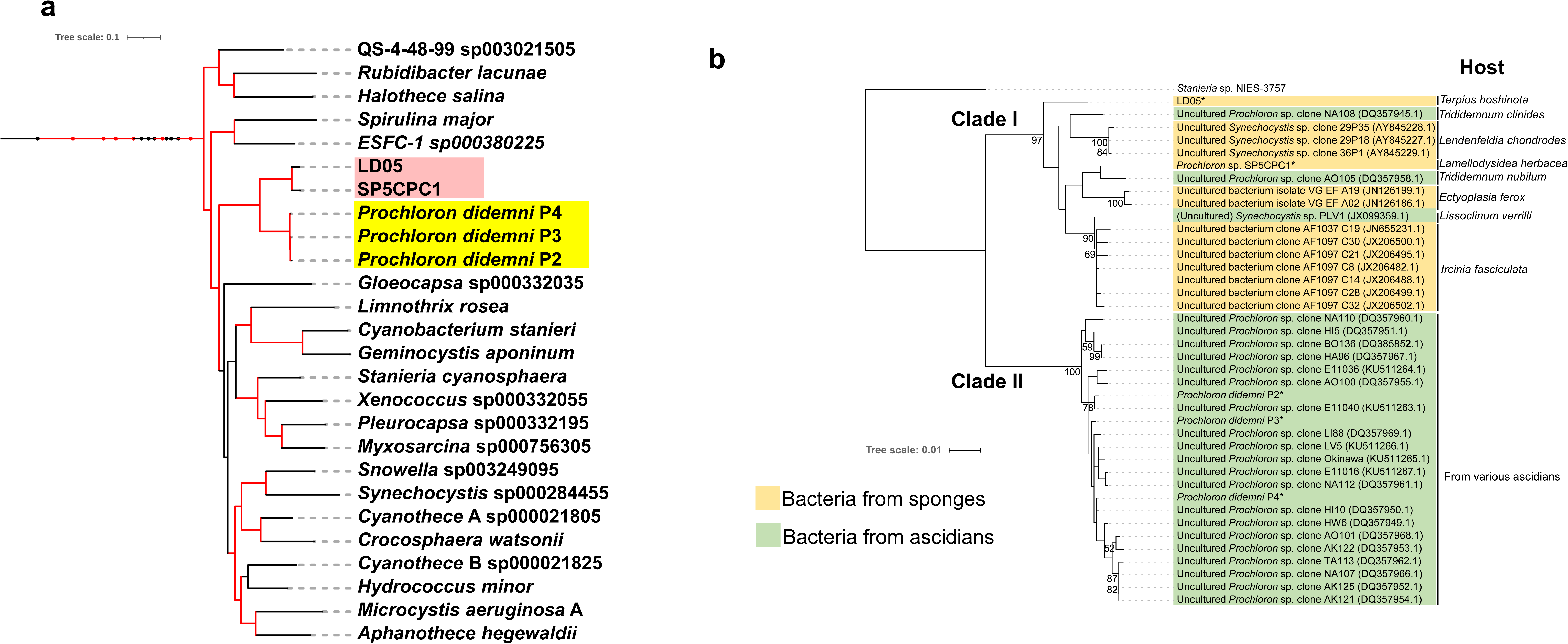
Molecular phylogenetic analyses of *Prochloron* and *Prochloron*-related bacteria. (a) Pruned phylogenetic tree based on the concatenation of 120 single-copy gene protein sequences. The complete phylogenetic tree including other cyanobacterial genera can be found in Supplementary figure 5. The branches with Ultrafast bootstrap (UFBoot) value >95% are highlighted with the red. The *Prochloron* genomes are labeled with yellow, and LD05 and SP5CPC1 are labeled with pink. *Vampirovibrio chlorellavorus* C was used as the outgroup. (b) A phylogenetic tree was constructed using the 16S rRNA gene by the maximum-likelihood method with 1,000 bootstraps. The tree included 16S rRNA gene sequences from SP5CPC1, LD05, *Prochloron* genomes—including *P. didemni* P2, *P. didemni* P3, and *P. didemni* P4—and other related 16S rRNA gene sequences identified by PCR cloning from environmental samples. The 16S rRNA gene sequence of *Stanieria* sp. NIES-3757 genome was used as an outgroup. The scare bar represents the number of changes per nucleotide. Asterisk represents the genomes are available.

To characterize the functional differences between the bacteria in the two clades, LD05, SP5CPC1, and *P*. *didemni* genomes were annotated with KEGG orthologs and clusters of orthologous groups (COGs). Complete-linkage clustering of COG category abundances demonstrated that the LD05 and SP5CPC1 were more functionally similar to *Microcystis* groups than to *P*. *didemni* (Fig. S6).

Two hallmarks of *Prochloron* are the presence of chlorophyll *b* and absence of phycobilins in the cell (30). However, a previous study showed that the symbiotic cyanobacteria in *T*. *hoshinota* had phycobilins based on microspectrophotometry results (15), which is consistent with our finding that phycobilins synthase and the phycobiliproteins genes were present in LD05 and SP5CPC1 genomes. The absence of the chlorophyll *b* synthase gene in LD05 and SP5CPC1 genomes implied that the bacteria lacked chlorophyll *b*. In addition, the failure to identify chlorophyll *b* in the ascidian *Trididemnum nubilum* holobiont, which the PCR clone AO15 (DQ357958.1) in Clade I was recovered from, implied that the bacterium also lacks chlorophyll *b*. In accordance with the genomic analyses, our pigment analysis by LC-QTOF-MS also revealed the presence of chlorophyll *a* and absence of chlorophyll *b* in *T*. *hoshinota* (Fig. 5). These results indicated that the bacteria in Clade I had different light-harvesting systems from *Prochloron* in Clade II.

**Figure 5.**
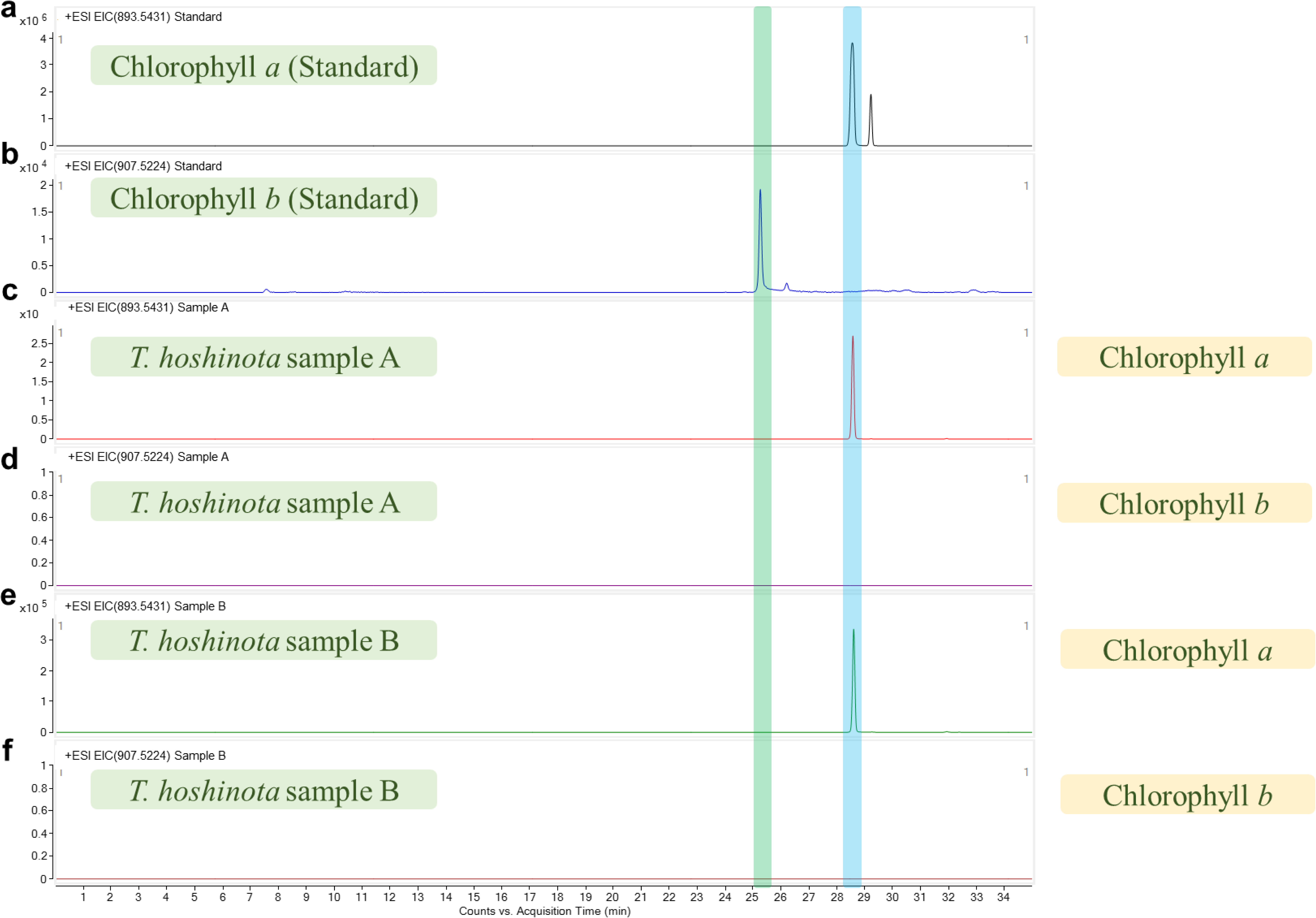
Extracted ion-chromatograms (EICs) of UPLC-QTOF-MS of the extracts of *T. hoshinota.* Chlorophyll *a* analytical standard (a) and chlorophyll *b* analytical standard (b) were used in the analysis. The analysis included two *T. hoshinota* samples. EIC of MS spectra within the m/z value of 893.543 (a, c, e) and 907.522 (b, d, f). a: chlorophyll *a* peak. b: chlorophyll *b* peak.

In summary, the results indicate that the LD05 and SP5CPC1 were derived from a common ancestor, phylogenetically different from the ancestor of *Prochloron*. More importantly, the bacteria have photosynthetic machinery distinct from that of *Prochloron*. The existence of phycobiliproteins and lack of chlorophyll *b*, inconsistent with the iconic definition of *Prochloron.* Therefore, we propose that the bacteria in Clades I and II be classified into two different genera and classify the species in Clade I as a novel genus “Paraprochloron”. Moreover, we classified LD05 as a novel species named *Candidatus* Paraprochloron terpiosi LD05 [Etymology: Gr. pref. *para*-, beside, alongside of; N.L. neut. n. *Prochloron*, a bacterial generic name; N.L. neut. n. *Paraprochloron*, a genus adjacent to *Prochloron*. N.L. gen. n. *terpiosi*, of *Terpios* a zoological scientific genus name.].

### Metabolic features of the novel species *Ca.* Pp. terpiosi LD05

*Ca*. Pp. terpiosi LD05 possessed nearly all the genes required for photosynthesis, carbon fixation, the tricarboxylic acid cycle (TCA), and glycolysis (Fig. 6). In addition, genes related to sucrose metabolism—e.g., sucrose-6-phosphatase, sucrose synthase, and sucrose phosphorylase—were identified. These genes were also present in the SP5CPC1 genome, but not in *P*. *didemni* genomes (Table S4). For light-harvesting systems, like typical cyanobacteria, the genes encoding phycobiliproteins—including phycoerythrin, phycocyanin, and allophycocyanin—were recovered, except for *cpcC*, *cpcD*, and *cpeE* (Table S4). Phycobilin synthase genes were also detected in the genomes. On the other hand, the genes encoding chlorophyll *a* synthesis were present, but the gene for chlorophyll *b* synthase was absent.

**Figure 6.**
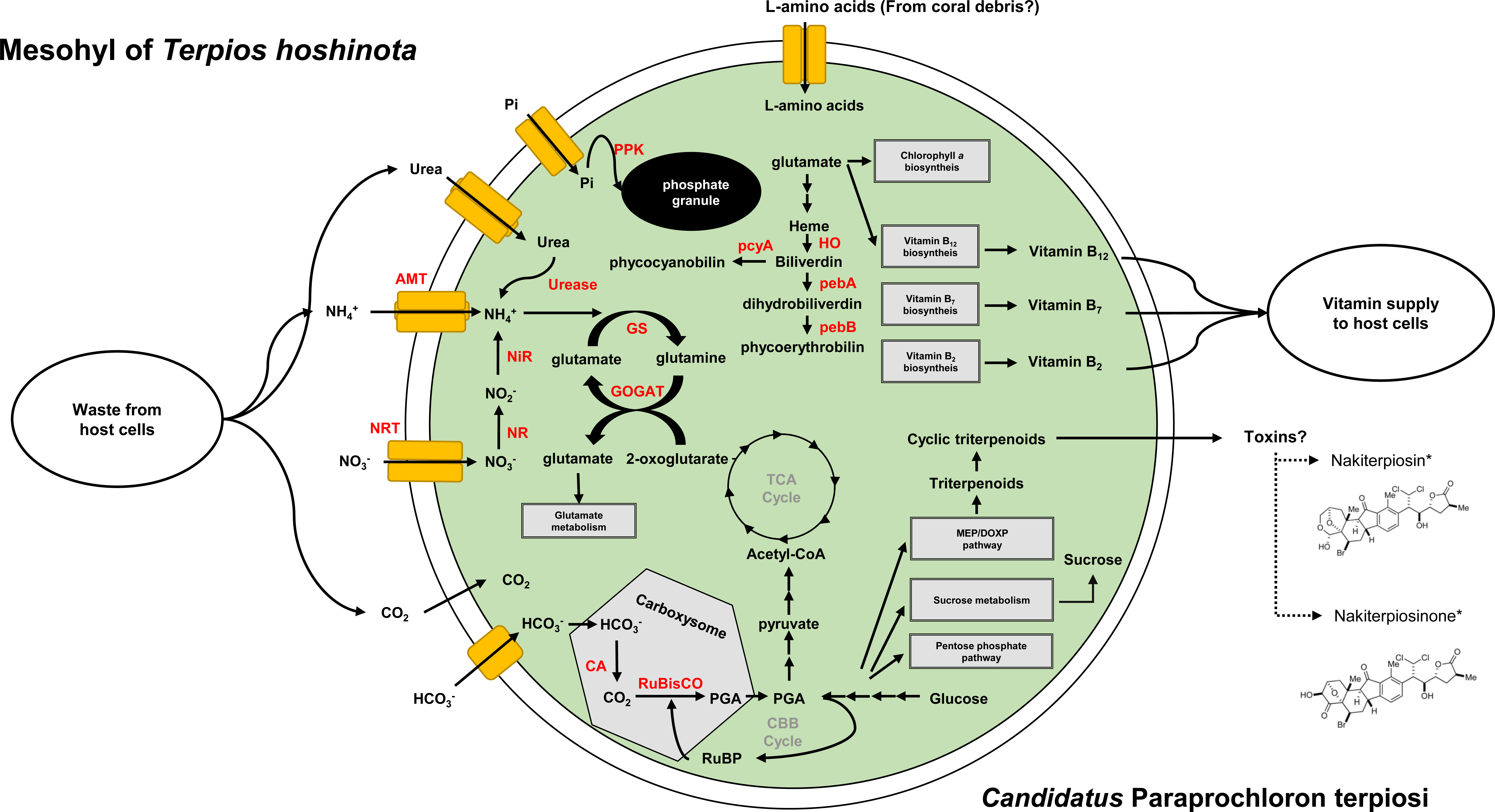
Metabolic potentials and putative nutrient cycling between *T*. *hoshinota* and *Ca*. Pp. terpiosi LD05. The bacterium may recycle nitrogen waste from host cells and provide them with vitamins. The bacterium may also produce secondary metabolites as toxins to help the sponge weaken coral tissues and facilitate overgrowth. On the other hand, the bacterium may also store phosphate as an energy reservoir for the holobiont to prepare for the competition with coral. Abbreviation: amt, ammonia transporter; NRT, nitrate transporter; NR, nitrate reductase; NiR, ferredoxin--nitrite reductase; CA, carbonic anhydrase; PPK, polyphosphate kinase; GS, glutamine synthetase; GOGAT, glutamine oxoglutarate aminotransferase; RuBisCo, ribulose-1,5-bisphosphate carboxylase/oxygenase; pcyA, phycocyanobilin:ferredoxin oxidoreductase; HO, heme oxygenase; pebA, dihydrobiliverdin:ferredoxin oxidoreductase; pebB, phycoerythrobilin:ferredoxin oxidoreductase. *The structures of nakiterpiosin and nakiterpiosinone are depicted based on Shuanhu Gao et al (2012).

Certain symbiotic cyanobacteria contain nitrogen fixation machinery and provide their host with nitrogen sources (31). However, the *Ca*. Pp. terpiosi LD05 and SP5CPC1 lacked nitrogen fixation genes. *Ca*. Pp. terpiosi LD05 had genes for nitrate uptake and assimilatory nitrate reduction pathways to convert nitrate into ammonium (Table S4). The glutamine synthetase/glutamate synthase (GS/GOGAT) pathway was also found in the genome, indicating that the ammonium derived from extracellular nitrate can be incorporated into amino acids by this pathway. Additionally, the genome also carried the complete gene set for the urea transporter and urease. The bacterium may take in the urea from hosts and convert it into ammonium. More importantly, the general L-amino acid transporters can be identified in the genomes, which cannot be observed in *Prochloron* genomes. Hence, the bacteria may have multiple methods for obtaining nitrogen resources from their host and environment.

Many symbiotic bacteria in sponges have been shown to provide their hosts with cofactors and vitamins such as vitamin B_12_ (cobalamin) (Table S4), which can only be synthesized by prokaryotic organisms (22). The *Ca*. Pp. terpiosi LD05 and SP5CPC1 genome contained nearly complete gene sets for the synthesis pathways of vitamin B_1_ (thiamine), vitamin B_2_ (riboflavin), vitamin B_7_ (biotin), and vitamin B_12_. The transporter for cobalt, a key constituent of vitamin B_12_, was also be identified. In contrast, the *Prochloron* genomes lacked a cobalt transporter and many genes for vitamin B_12_ synthesis.

Previous studies have discovered a variety of secondary metabolites from the *T*. *hoshinota* holobiont (32); these metabolites may be produced by symbiotic bacteria (33). Our analysis revealed the existence of four terpene synthesis clusters in *Ca*. Pp. terpiosi LD05, as well as SP5CPC1. Furthermore, the species also contains a gene encoding squalene cyclase, suggesting that the bacterium can produce cyclic triterpenes. Other secondary metabolite biosynthetic gene clusters were also recovered. The gene clusters present in *Ca*. Pp. terpiosi LD05 included butyrolactone, linear azol(in)e-containing peptides, and non-ribosomal peptide synthetase clusters (Fig. 3).

### Symbiotic signatures and unique genomic features of *Ca.* Pp. terpiosi LD05

The complete genome of *Ca*. Pp. terpiosi LD05 enables us to investigate the genome feature in a precise and comprehensive manner. When we compared the genomes with other phylogenetically close cyanobacteria, we found that the *Ca*. Pp. terpiosi LD05 and SP5CPC1 lacked *dnaA* gene, which encodes protein for replication initiation factor responsible for DNA unwinding at *oriC* (34). Previous studies mentioned that many of symbiotic cyanobacteria lacked *dnaA* gene and adopt DnaA-independent replication mechanisms, which may lead to multiple replicative origins and polyploidy (35, 36). Another interesting feature of *Ca*. Pp. terpiosi LD05 we found was that the bacteria had 50 copies of RNA-directed DNA polymerase genes, which was not observed in other phylogenetically close cyanobacteria. However, the function of this feature needs to be determined.

Many symbiotic bacteria in sponge carry eukaryotic-like proteins (ELPs). The ELPs— e.g., leucine-rich repeat proteins (LRRs), ankyrin repeat proteins (ARPs), WD-40 containing proteins (WD-40), and tetratricopeptide repeat proteins (TPRs)—may be used to interact with hosts during various biological processes. The genomic analysis demonstrated that *Ca*. Pp. terpiosi LD05 carried these ELPs. However, some closely related free-living cyanobacteria also possess ELPs. On the other hand, the ELPs were highly enriched across *Prochloron* genomes.

## Discussion

*T. hoshinota* is one of the most important biological threats to coral in the Indo-Pacific region. Its associated cyanobacteria and other symbiotic bacteria are essential for maintaining the sponge’s function. Nevertheless, the present study is the first to explore the composition and role of *T. hoshinota*’s microbiome. Our unprecedented large-scale survey of the bacterial community in *T*. *hoshinota* based on 16S amplicon sequencing of samples from various regions across the western Pacific Ocean, Indian Ocean, and South China Sea enabled us to characterize the sponge’s core microbiome.

We identified a bacterium, closely related to *Prochloron*, that was dominant across our sponge samples. The prevalence and predominance of this bacterium in *T*. *hoshinota* highlight the strong relationship between bacteria and their hosts. Combining Nanopore and Illumina sequencing, we recovered the complete bacterial genome with high accuracy, followed by phylogenetic and functional analyses, demonstrated that this cyanobacterium has a distinct phylogenetic affiliation and genomic and metabolic characteristics from *Prochloron,* indicating that it belongs to a novel genus. We proposed a new genus, “Paraprochloron”, and refined the membership of *Prochloron*. Lastly, a functional analysis was performed to understand the lifestyle of this novel bacterium in the *T. hoshinota* holobiont. This study provides insights into the role of the cyanobacterium in *T. hoshinota*, which will facilitate future studies to investigate the ecosystem inside *T. hoshinota*, the causes of its outbreaks, and its invasion mechanism. Recovery of the complete bacterial genome will enable in-depth molecular approaches for identification of ecological functions of this cyanobacterium, and the sponge-cyanobacteria interactions

### Biogeographical variation in the *T*. *hoshinota* microbiome and its core microbial **members**

The microbial community in sponges can be shaped by several environmental factors—e.g., light, nutrients, temperature, and pH—and may change in response to stress (23, 24, 37, 38). Environmental changes may force a sponge holobiont to alter its microbial community to acclimatize (39). On the other hand, host-related factors can also be important for determining community composition. For example, previous studies have shown that sponges have several immune receptors, such as NOD-like and Toll-like receptors (13). These receptors enable sponges to discriminate between symbionts and prey. In order to live inside a sponge, microbes have to evade host immunity and phagocytosis (40, 41). Furthermore, metabolites and nutrients provided by hosts also contribute to the community structure (42). Some sponges show high stability in their microbiome due to strict selective pressures from their host. For instance, *Ircinia* and *Hexadella* sponges exhibit host-specific and stability in their associated bacterial communities, despite large geographic distances between sampling sites (24, 43). In contrast, certain sponges, such as *Petrosia ficiformis,* harbor biogeographical-dependent bacterial compositions (44).

Our survey of three oceans enabled us to determine whether bacterial communities vary across different biogeographical backgrounds and experience strong selective pressure from their host. Our dissimilarity analysis of the microbial composition of *T. hoshinota* showed a weak correlation between sample location and bacterial composition. The differences in microbial composition among *T. hoshinota* samples from different locations may result from local acclimatization. Samples from the same oceanic regions that share communities may equip their holobionts with similar metabolic functions to cope with stress or increase fitness in certain environments.

However, despite the biogeographical variation, we still identified core bacterial members that were present in all samples (Fig. S3). This core bacterial community comprised only a few genera, but accounted for ∼80% of the community across all samples, indicating that it is vital to the holobiont. Host-related factors may help keep this core microbiome stable because its members carry out core functions of the holobiont (23). The core members may contain the metabolic capability to utilize nutrients from the sponge host’s environment and play important roles in nutrient exchanges, such as sulfur, carbon, and nitrogen cycling. This core group may be also responsible for defending against predators and protecting the sponge symbionts from toxins and pathogens (23).

The most dominant OTU in the core microbiome was closely related to *Prochloron*, a genus of symbiotic cyanobacteria found in various ascidians. The biogeographically- independent prevalence and predominance of this *Prochloron*-related bacterium in *T. hoshinota* suggest that it has a crucial role in the holobiont.

### *Candidatus* Paraprochloron terpiosi gen. nov., sp. nov., a *Prochloron*-related bacterium prevalent in *T. hoshinota*

Although the 16S rRNA gene of the *Prochloron*-related bacterium in *T. hoshinota* is phylogenetically close to that of *Prochloron*, deeper genomic, phylogenetic, and functional analyses, combined with the recent recovery of another symbiont *Prochloron*-related bacterial genome in the sponge, revealed that the two cyanobacteria are distinct, especially in terms of their pigment contents and phylogenetic divergence. Moreover, the pigment features in these bacteria inconsistent with the definition of *Prochloron*. Hence, a new genus, “Paraprochloron”, was formally erected to distinguish these two groups.

Our 16S rRNA metagenomic analysis revealed that *Candidatus* Paraprochloron terpiosi LD05 is the most dominant cyanobacterium within *T*. *hoshinota* specimens collected across three different oceans. Why does the dominant cyanobacterium species remain identical in *T. hoshinota* samples across different oceans? The reason remains unclear. Recent studies suggest that *T. hoshinota* larvae, which carry vertically transmitted cyanobacteria, may have a short dispersal distances because they are more dense than the water, leading them to settle rapidly after leaving their mother sponge (16–18). In this circumstance, the symbiotic *Ca*. Pp. terpiosi LD05 from various locations might accumulate genetic differences to adapt to local environments. The absence of evident speciation in our study may be the result of the tight and stable symbiotic interactions between *Ca*. Pp. terpiosi LD05 and *T. hoshinota* restricting genetic changes in the bacterium. Another possibility is that the species becomes broadly dispersed through some other mechanism, such as via ocean currents or transportation via vessels; this would enable *T. hoshinota* larvae to spread quickly across the ocean with few genetic alterations. Evidence of recent *T. hoshinota* outbreaks support this latter scenario (2).

### Comparison between “Paraprochloron” and *Prochloron*

Comparative genomics between “Paraprochloron” and *Prochloron* enables us to infer their evolutionary histories and respective relationship to their host. Previous studies have shown that symbiotic bacteria usually have reduced genomes because certain genes erode as symbiosis develops (45, 46). Furthermore, a model of symbiont evolution has proposed that during the evolutionary history of symbiosis large-scale pseudogenization can happen during transitional events, such as a strict host association or vertical transmission, leading to a sudden drop in coding density (46). Eventually, the coding density gradually bounces back due to deletion bias.

Comparing *Prochloron* and “Paraprochloron”, we found that “Paraprochloron” had a smaller genome but similar coding density to other phylogenetically closely related cyanobacteria (Table S4). *Prochloron*, on the other hand, had a similar genome size but lower coding density. This difference may indicate that the transition toward the host-restricted lifestyles occurred more recently in *Prochloron* than “Paraprochloron”. The hypothesis can be supported by our observation that *Prochloron* harbored highly-enriched transposase genes as the elevation of mobile genetic element quantities, such as transposons and insertion sequences, is thought to be associated with the recent transition to a host-restricted symbiotic lifestyle (47).

Along with its unusual pigments, another hallmark of *Prochloron* is that is produces patellamides, cytotoxic cyclic peptides. However, the synthesis gene cluster cannot be found in *Ca*. Pp. terpiosi LD05 or SP5CPC1. On the contrary, the latter two genomes have four terpene synthesis gene clusters and contain genes that encode the protein that circulates terpene, but *Prochloron* genomes only harbor two terpene synthesis gene clusters. Hence, “Paraprochloron” and *Prochloron* may produce distinct secondary metabolites to increase the fitness of themselves or their entire holobiont.

Certain cyanobacteria contain genes for sucrose metabolism. Though the role of sucrose in cyanobacteria remains understudied, several articles have shown that sucrose can be utilized as a compatible solute, serve as signal molecule, or be used for glycogen synthesis (48–51). Interestingly, our genomic analysis found that the genes involved in sucrose metabolism were present in *Ca*. Pp. terpiosi LD05 and SP5CPC1 but absent in *Prochloron* (Table S4). We hypothesize that “Paraprochloron” can use sucrose as an osmolyte to cope with osmotic stress; this is supported by a previous finding that the genes related to osmotic stress are enriched in sponge-associated bacterial genomes (22). On the other hand, we also found that *Ca*. Pp. terpiosi LD05 and SP5CPC1 contained osmoprotectant transporter genes, which were not identified in *Prochloron* genomes (Table S4). These results indicate that the “Paraprochloron” species may live in environments with higher osmotic stress, like places with high osmolarity or fluctuations in osmolarity, compared to *Prochloron*. Alternatively, “Paraprochloron” and *Prochloron* may use different strategies to deal with osmotic stress.

Another observation that drew our attention is the absence of *dnaA* gene in *Ca*. Pp. terpiosi LD05 and SP5CPC1. DnaA is required for the initiation of DNA replication at *oriC* (34). Some bacterial symbionts do not possess *dnaA* gene (36, 52, 53). Like mitochondria and chloroplasts, certain *dnaA*-lacking symbionts had multiple copies of the same genomes, leading to the increase in genome copy number in a single cell (35). The evolution of DnaA-independent replication was most studies in cyanobacteria. Many symbiotic cyanobacteria were found lacking *dnaA* gene. A recent study found that the deletion of the *dnaA* gene was not lethal in some free-living cyanobacteria, and deletion of the gene increased cell viability in stationary phase (36). Another study also indicated the cyanobacteria lost DnaA dependency before they become symbiosis, and the loss will drive the free-live bacteria to became symbiosis (35, 36). However, in our analysis, *Prochloron* has *dnaA,* but “Paraprochloron” does not, indicating the loss of *dnaA* happened after the two symbionts separate from the same ancestor. It implies that the symbiosis may also drive the bacteria to lose *dnaA*. However, the function of *dnaA*-independent replication in symbiosis still requires further experimentation.

The eukaryotic-like proteins (ELPs) are highly-enriched in *Prochloron* but not “Paraprochloron”. ELPs can be used to perform protein-protein interactions, phagocytosis evasion, host cell binding, and interference with the host ubiquitin system (54). ELPs are also thought to be necessary for sponge symbionts (8). The differences in ELP quantities between *Prochloron* and “Paraprochloron” may indicate that “Paraprochloron” uses an unidentified mechanism to interact with their hosts. Another possibility is that the long-term symbiotic history of “Paraprochloron” propelled bacteria to retain their essential ELPs and reduce unnecessary ones.

### Symbiotic interactions between *Ca*. Pp. terpiosi and *T. hoshinota*

Sponges harbor complex bacterial communities (13), and the symbiotic bacteria play vital and diverse roles in the physiology and ecology of sponges. For instance, bacteria can provide their hosts with various nutrients and recycle waste. Moreover, symbiotic microorganisms can produce diverse secondary metabolites to compete with other holobionts or defend from invasion by predators. The predominance of *Ca*. Pp. terpiosi LD05 highlights its role in the *T. hoshinota* and the genome of *Ca*. Pp. terpiosi LD05 enables us to understand its metabolic potential, the symbiotic relationship, and even how the holobiont overgrows corals.

Many sponges are mixotrophs and certain sponges can obtain carbon resources from cyanobacteria (55). Some sponges even acquire more than 50% of their energy demand from the symbiotic cyanobacteria in the form of photosynthates (56). Photosynthesis was observed in the *T. hoshinota* holobiont, and its efficiency increased when the coral-killing holobiont came into contact with coral; this mechanism may help sponges overgrow the coral (57). *Ca*. Pp. terpiosi LD05 is the only cyanobacterium in the core microbiome, and it has been found to have the genes needed for photosynthesis. Hence, the bacterium may be an important carbon source for *T. hoshinota* and facilitate competition with coral.

Ammonia secreted by sponges may serve as a nitrogen source for symbiotic bacteria. Our analysis identified ammonium transporters and the GS-GOGAT pathway in *Ca*. Pp. terpiosi LD05. Hence, the bacterium may recycle the ammonium from host. On the other hand, the bacterium harbors the urea transporter and urease. Therefore, urea can be used as an alternative nitrogen source. A previous study showed that the level of free amino acids in *T. hoshinota*-living cyanobacteria increased when the holobiont encountered coral. Our finding of an amino acid transporter in *Ca*. Pp. Terpiosi LD05 suggests that the bacterium may benefit from “coral killing” by leaching amino acids or ammonium that are released as the coral disintegrates.

Some microbes that live in sponges can store polyphosphate granules, which can account for 25–40% of the total phosphate content in sponge tissue (58, 59). These polyphosphate granules can be used for energy storage and may enable the holobiont to withstand periods of phosphate starvation. We found genes encoding polyphosphate kinase. Hence, *Ca*. Pp. terpiosi LD05 may play a pivotal role in phosphate storage and cycling in the *T. hoshinota* holobiont. A previous study demonstrated that the phosphate concentration in seawater was highly correlated with *T. hoshinota* overgrowth (60), highlighting the importance of phosphate in the spread of the *T. hoshinota* holobiont. Another study also proposed that *T. hoshinota* accumulates nutrients to compete with corals (8). We propose that *Ca*. Pp. terpiosi LD05 helps *T*. *hoshinota* outcompete corals by serving as a carbon and phosphate storage. However, how the phosphate concentration is correlated with *T. hoshinota* cover is unclear, as are details about phosphate cycling among the host, *Ca*. Pp. terpiosi LD05, and other symbiotic microbes.

Animals cannot synthesize essential vitamins, so symbiotic microorganisms are thought to be important sources of essential vitamins for sponges. Our analysis of the *Ca*. Pp. terpiosi LD05 genome identified the biosynthesis pathway for vitamin B_1_, vitamin B_7_, and vitamin B_12_. Thus, *Ca*. Pp. terpiosi LD05 may help maintain *T. hoshinota*’s health by providing the holobiont with vitamins.

One of the leading questions in the study of *T. hoshinota* is how it kills corals. Several mechanisms have been proposed. One argues that the sponge produces cytotoxic allelochemicals to damage coral cells (61). Another suggests that the sponge overgrows corals and competes with them for nutrients (8). Finally, the sponge may cause the coral’s microbial composition to change, causing it to malfunction (62). These hypotheses are not mutually exclusive. For the first, several cytotoxic compounds, including nakiterpiosin, nakiterpiosinon, and terpiodiene, have been isolated from *T. hoshinota* holobionts (63, 64). Nakiterpiosin, and nakiterpiosinon are C-nor-D-homosteroids. Previous studies showed that cyanobacteria can produce sterols by the cyclization of squalene, a triterpene (65). In the *Ca*. Pp. terpiosi LD05 genome, the synthesis pathway of squalene and squalene cyclases were identified. Hence, the bacterium may be responsible for the production of the toxins. These toxins may facilitate overgrowing corals by damaging coral tissue directly or weakening the coral’s defenses. In the future, the products of these gene clusters can be confirmed by molecular approaches.

### Conclusions

This study makes several fundamental discoveries about the bacterial community associated with *T. hoshinota*. The core microbiome of *T. hoshinota* includes a *Prochloron-*like bacterium, as well as *Endozoicomonas*, *Pseudospirillum*, SAR116, *Magnetospira*, and *Ruegeria*. Our finding that the dominant cyanobacteria in *T. hoshinota* is the same throughout the western Pacific Ocean, South China Sea, and Indian Ocean suggests that this particular cyanobacterium is the obligate symbiont of the sponge. The complete genome of this cyanobacterium was recovered, and the phylogenetic and genomic analyses revealed that the genome should be classified as a novel species under a new genus “Paraprochloron”, close to *Prochloron*. The complete genome enables us to understand the carbon and nitrogen metabolism of the bacterium, and the putative symbiotic interaction between the bacterium and *T*. *hoshinota*. By identifying the core microbiome and revealing the metabolic potential of the dominant cyanobacterium in *T*. *hoshinota*, our conclusions will help future studies explore the detailed ecosystem inside the holobiont, unveil how *Ca*. Pp. terpiosi LD05 contributes to the invasion and coral-killing nature of *T*. *hoshinota*, and determine the cause of the outbreaks in the future. Also, the complete genome we reconstructed will help us to extend our knowledge of cyanobacteria evolution and the functional diversity of symbiotic cyanobacteria.

## Materials and methods

### Sample collection

Sixty-one sponge samples, summarized in Supplementary table 1, were collected from 15 coastal sites in the western Pacific Ocean, South China Sea, and Indian Ocean. The collection sites in the western Pacific Ocean included Okinoerabu (JPOKE), Bise (JPBS), Miyako (JPMYK), and Ishigaki (JPISG) in Japan, Lyudao (TWLD), Lanyu (TWLY), and Kenting (TWKT) in Taiwan, Guam (USGU) in the United State, and the Great Barrier Reef (AUGBR) in Australia. The collection sites in the South China Sea included Yongxing (CNYX) and Taiping (TWTP) in Taiwan, Redang (MYRD) in Malaysia, and Pari (IDPR). in Indonesia. Indian Ocean samples were collected from Faafu Atoll (MVFF) in the Maldives. Before DNA isolation, sponge tissue was collected with tweezer underwater and placed in Falcon 50 mL conical centrifuge tubes. Samples were then washed with 1 ml 1X TE buffer.

### DNA extraction

For the sponge samples from JPBS, JPISG, JPMYK, JPOKE, TWLD, TWLY, and USGU, total DNA was extracted using a DNeasy Blood and Tissue kit (QIAGEN, Hilden, Germany) according to manufacturer’s protocol. For the other samples, total DNA was extracted by a modified CTAB method described in our previous study (14). After DNA isolation, the DNA were stored at -80°C until subsequent experiments.

### 16S rRNA gene amplification and multiplex tag sequencing

High throughput sequencing of the 16S rRNA hypervariable V6–V8 region was used to characterize bacterial community diversity and composition. V6–V8 sequences were amplified by PCR with forward primer 5µ-AACGCGAAGAACCTTAC-3µand reverse primer 5µ-GACGGGCGGTGWGTRCA-3µ (66). PCR mixtures contained 33.5 µL of sterilized distilled water, 0.5 µL of 5 U TaKaRa Ex Taq (Takara Bio, Otsu, Japan), 5 µL of 10X Ex Taq buffer, 4 µL of 10 mM dNTP, 1 µL of each primer at a concentration of 10 µM, and 5 µL template DNA in a total volume of 50 µL. PCR was programmed with an initial step of 94°C for 5 min, 30 cycles of 94°C for 30 s, 52°C for 20 s, and 72°C for 30 s, followed by a final step of 72°C for 10 min. PCR was performed again to add barcodes to the amplicons. Each primer tag was designed with four extra nucleotides at the 5’ end of both primers. Unique tags were used for PCR barcoding to label each sample in the study. PCR conditions were the same as that for the V6–V8 amplification, except that the number of reaction cycles was reduced to five. The products were purified by 1.5% agarose electrophoresis and QIAEX II Agarose Gel Extraction kit (QIAGEN, Hilden, Germany) according to manufacturer’s instruction. The quality of the purified products was assessed using a Nanodrop spectrophotometer (Thermo Scientific, Vantaa, Finland).

### Illumina sequencing and community analysis

DNA concentration was measured by a Quant-iT™ assay (Thermo Fisher Scientific, USA). Equal pools of DNA were sent to Yourgene Bioscience (Taipei, Taiwan) and sequenced using the MiSeq platform (Illumina, USA). Short and low-quality reads were filtered by Mothur v.1.38.1 (67) to retain reads with an average quality score > 27 and lengths of 365–451 base pairs (bp). Reads with homopolymer > 8 bp were excluded, and chimera sequences from all samples were removed by USEARCH v8.1.1861 (68). OTUs were determined and classified using the UPARSE pipeline (69) with reference to the Silva v128 database (70). OTUs were defined using a 97% similarity threshold. OTUs assigned to Archaea, Eukaryota, chloroplast, and mitochondria were removed.

Alpha and beta diversity indices were determined by QIIME 2 v2020.6 (71). Alpha diversity indices were calculated by Shannon diversity, Chao1 richness estimator, and Faith’s phylogenetic diversity. Dissimilarities in microbial community compositions among samples were computed using Bray–Curtis distance dissimilarity, unweighted UniFrac, and weighted UniFrac. The results were visualized by principal component analysis (PCA) and dendrogram.

### Electron microscopy

*T*. *hoshinota* samples at the sponge-coral border were collected by transmission electron microscopy (TEM), and *T*. *hoshinota* was observed on rubber by scanning electron microscopy. Samples were prepared as described in our previous study (14). They were fixed in 0.1 M phosphate buffer with 2.5% glutaraldehyde and 4% paraformaldehyde. The samples were then washed with 0.1 M phosphate buffer for 15 min three times, then immersed in 0.1 M phosphate buffer with 1% osmium tetroxide for 4 hr. After being washed again three times, the samples for TEM were sequentially dehydrated in acetone at 30%, 50%, 70%, 85%, 95%, and 100% for 20 min each time. After being dehydrated, the samples were embedded in Spurr’s resin and sectioned. The sections were stained with 5% uranyl acetate in methanone and 0.5% lead citrate. The stained samples were observed under a Philips CM-100 TEM. Samples for SEM were sequentially dehydrated in 30% for 1 hr, 50% for 1 hr 70% for 1 hr, 85% for 2 hr, 95% for 2 hr, 100% for 2 hr, and 100% for 12 hr. The samples were dried in a Hitachi HCP-2, then gold coated with a Hitachi E-1010 and observed under an FEI Quanta 200 SEM.

### Pigment analysis by UPLC**-**QTOF-MS

Approximately 2.5 g of wet *T*. *hoshinota* tissue from each sample was centrifuged at 7,000 rpm for 30 s to remove excess water. Sponge tissues were placed in mortars and ground with 4 ml of 100% acetone to homogenize the tissues. The homogeneous samples were transferred into a 5 ml centrifuge tube and centrifuged at 7000 rpm at room temperature for 30 s. The supernatant was transferred into a new 5-ml centrifuge tube. The samples were covered with aluminum foil paper for being shaded and stored at 4°C until pigment analysis.

Pigment content analysis was performed by UPLC-MS. The UPLC-MS system used the Agilent 1290 Infinity II ultra-performance liquid chromatography (UPLC) system (Agilent Technologies, Palo Alto, CA, USA) coupled online to the Dual AJS electrospray ionization (ESI) source of an Agilent 6545 quadrupole time-of-flight (Q-TOF) mass spectrometer (Agilent Technologies, Palo Alto, CA, USA). The samples were separated using a Kinetex XB-C18 column (2.6 μm, 4.6 × 100 mm, Phenomenex, Torrance, CA, USA) at 40°C. The chromatogram was acquired; mass spectral peaks were detected and their waveform processed using Agilent Qualitative Analysis 10.0 software (Agilent, USA).

### Metagenome sequencing and assembly

Nanopore and Illumina sequencing were used to recover metagenome-assembled genomes (MAGs) with high accuracy. For Illumina sequencing, *T*. *hoshinota* tissue was ground, and the samples were filtered through a 0.35 μm membrane. 1 x 10^8^ cyanobacterial cells were purified from the samples using the MoFlo XDP cell sorter based on fluorescence and forward scatter intensity. The cells were then retained on a 0.2-μm cellulose acetate membrane filter, and the DNA was extracted by CTAB method mentioned above. Purified DNA was sent to Yourgene Bioscience (Taipei, Taiwan) and sequenced on a MiSeq platform (Illumina, USA).

For Nanopore sequencing. *T*. *hoshinota* larvae were collected into a Falcon 50 mL conical centrifuge tube with a dropper underwater. The larvae were then selected into a Petri dish using a microscope to remove contamination, and the larvae were then fixed with absolute ethanol and stored at -20°C until DNA extraction. The total DNA was extracted by a modified CTAB method mentioned above. After DNA extraction, the DNA was sent to NGS Core at Biodiversity Research Center, Academia Sinica for Nanopore sequencing. To recover cyanobacterial genome from the metagenome, Nanopore reads were assembled by metaFlye with default settings (26). One contig was annotated as circular by metaFlye, and it was assigned as novel cyanobacterial species by GTDB-TK taxonomy annotation (72). Because Nanopore reads are error-prone, the genome was polished by Illumina reads in order to increase the accuracy of the genome. First, Illumina reads were trimmed and filtered by trimmomatic v0.39 with the following parameters: ILLUMINACLIP:TruSeq3-PE-2.fa:2:30:10:3: TRUE LEADING:10 TRAILING:10 SLIDINGWINDOW:5:15 MINLEN:50 (73). The processed reads were mapped to the cyanobacterial contig, and the contig was then polished with Pilon by the mapping result (74).

### Genome annotation

The genome of *Prochloron*, “Paraprochloron”, and phylogenetically close cyanobacteria were annotated using Prokka v1.13.7 with the ‘usegenus’ and ‘rfam’ options (75). The genomes were also annotated with KEGG functional orthologs (KO numbers) and COGs (76, 77). To annotate the KO numbers, protein sequences predicted by Prodigal were blasted against the KEGG database using BlastKoala and enrichM (78–80). The K number annotation results were then used to reconstruct the transporter systems and metabolic pathways using KEGG mapper (81). To annotate COGs, the predicted protein sequences were searched against the COG database by PSI-BLAST with an e-value threshold of 10^-5^ (82). In addition, the transporter proteins were identified by searching the putative protein sequences against TransportDB 2.0 (August 2019) using BLASTp (83).

### Phylogenetic analysis

To reconstruct a phylogenetic tree of the 16S rRNA genes, the sequences with high identities to the 16S rRNA gene in LD05 genome were retrieved by searching the NCBI nt database using BLASTn (82, 84). The sequences were aligned by MUSCLE on MEGA7 (85, 86). A tree was then reconstructed using IQ-TREE v2.03 using automatic model selection and 1,000 bootstraps (87). The tree was visualized with iTOL v4 (88).

Genomes for each cyanobacteria species in GTDB database were downloaded and 120 single-copy gene protein sequences were used to reconstruct a tree (89). Marker gene protein sequences were identified, aligned, and concatenated by GTDB-TK v1.3.0 (72). A phylogenomic tree was then built using IQ-TREE v2.03 with automatic model selection and 1,000 Ultrafast bootstraps. The tree was visualized with iTOL v4 (87, 88).

### Availability of data and materials

All the raw data and genome were submitted under BioProject ID: PRJNA665642.

## Acknowledgements

Y.H.C acknowledges the Taiwan International Graduate Program (TIGP) for its fellowship towards his graduate studies. We thank Noah Last of Draft Editing for his English language editing, Dr. Ker-yea Soong and Shih-Shou Fang for collecting *T*. *hoshinota* samples, and the Green Island Marine Research Station for supporting the research. We thank the electron microscope and flow cytometry divisions of IPMB, Academia Sinica for their technical support and Chih-Yu Lin and Gong-Min Lin in the Metabolomics Core Facility, Agricultural Biotechnology Research Center, Academia Sinica for the technical support and for performing the UPLC-MS/MS analysis and processing data.

## Competing interests

The authors declare that they have no competing interests.

## Funding

This work was funded by Biodiversity Research Center, Academia Sinica and the National Science Council, Taiwan (NSC98-2321-B-001-025-MY3678106).

## Authors’ contributions

YHC, HJC, and SLT designed the study and prepared the manuscript. YHC and HJC analyzed and interpreted the data. HJC, CYY, JHS, DZH, PWC, MHL, and WSC collected and sequenced the samples. HJC, CYY, and JHS observed the electron microscopy. HJC, THS, SHL, and CMY examined the pigments. CAC, JDR, EH, BHI, HH, PJS, CHJT, and HY assisted with the sampling.

All authors read and approved the manuscript.

---

**Table 1.** Basic statistics of the Candidatus Paraprochloron terpiosi LD05

## Supplemental Materials

**Figure S1**. Alpha diversity plots of each sample from different locations. (a) OTU richness estimated by Chao1. (b) Faith’s Phylogenetic Diversity. (c) Shannon’s evenness index box plot. The lines extending outside the boxes indicate variability outside the upper and lower quartiles.

**Figure S2.** PCA visualizations and dendrograms of beta diversity analysis. PCA visualizations (a–c) and dendrograms (d–f) of beta diversity analysis using the Bray– Curtis metric (a and d), unweighted (qualitative) UniFrac (b and e), and weighted (quantitative) UniFrac (c and f).

**Figure S3.** Taxonomic compositions of the *T*. *hoshinota* microbiome from different locations at the phylum (a) and genus (b) levels based on 16S rRNA amplicon sequencing analysis.

**Figure S4.** Matrix of average amino acid identity between cyanobacteria that are closely related to the LD05 genome. The dendrogram was generated using the neighbor joining method.

**Figure S5.** Complete phylogenetic tree of Figure 4a. The tree includes 120 cyanobacteria from different genera. The branches with Ultrafast bootstrap (UFBoot) value >95% are highlighted with the red. The *Prochloron* genomes are labeled with yellow, and LD05 and SP5CPC1 are labeled with pink. *Vampirovibrio chlorellavorus* C was used as the outgroup.

**Figure S6.** Heatmap representing the relative abundance of genes assigned to COG functional categories. The analysis included SP5CPC1, LD05, *Prochloron*, and the members in the adjacent clade. The dendrogram was draw using complete-linkage clustering.

**Table S1.** Summary of information on sampling and reads

**Table S2.** Proportion of different cyanobacterial genera in *T*. *hoshinota*-associated cyanobacteria

**Table S3.** Genomic features of LD05, *Prochloron*, and other phylogenetically-close cyanobacteria

**Table S4.** Comparison of metabolic feature in “Paraprochloron” and *Prochloron*

## Notes

### Competing Interest Statement

The authors have declared no competing interest.

## References

1. Rützler K, Muzik K. 1993. Terpios hoshinota, a new cyanobacteriosponge threatening Pacific reefs. Recent advances in systematics and ecology of sponges Scientia Marina.

2. Liao M-H, Tang S-L, Hsu C-M, Wen K-C, Wu H, Chen W-M, Wang J-T, Meng P-J, Twan W-H, Lu C-K. 2007. The" Black Disease" of Reef-Building Corals at Green Island, Taiwan-Outbreak of a Cyanobacteriosponge. Terpios hoshinota (Suberitidae; Hadromerida). ZOOLOGICAL STUDIES-TAIPEI- 46:520.

3. Fujii T, Keshavmurthy S, Zhou W, Hirose E, Chen C, Reimer J. 2011. Coral-killing cyanobacteriosponge (Terpios hoshinota) on the Great Barrier Reef. Coral Reefs 30:483–483.

4. de Voogd NJ, Cleary DFR, Dekker F. 2013. The coral-killing sponge Terpios hoshinota invades Indonesia. Coral Reefs 32:755–755.

5. Hoeksema BW, Waheed Z, de Voogd NJ. 2014. Partial mortality in corals overgrown by the sponge Terpios hoshinota at Tioman Island, Peninsular Malaysia (South China Sea). Bulletin of Marine Science 90:989–990.

6. Thinesh T, Jose PA, Hassan S, Selvan KM, Selvin J. 2015. Intrusion of coral-killing sponge (Terpios hoshinota) on the reef of Palk Bay. Current Science 109:1030–1032.

7. Montano S, Chou WH, Chen CA, Galli P, Reimer JD. 2015. First record of the coral- killing sponge Terpios hoshinota in the Maldives and Indian Ocean. Bulletin of Marine Science 91:97–98.

8. Wang JT, Chen YY, Meng PJ, Sune YH, Hsu CM, Wei KY, Chen CA. 2012. Diverse Interactions between Corals and the Coral-Killing Sponge, Terpios hoshinota (Suberitidae: Hadromerida). Zoological Studies 51:150–159.

9. Plucerrosario G. 1987. The Effect of Substratum on the Growth of Terpios, an Encrusting Sponge Which Kills Corals. Coral Reefs 5:197–200.

10. Yomogida M, Mizuyama M, Kubomura T, Reimer JD. 2017. Disappearance and Return of an Outbreak of the Coral-killing Cyanobacteriosponge Terpios hoshinota in Southern Japan. Zoological Studies 56.

11. Shi Q, Liu G, Yan HQ, Zhang HL. 2012. Black Disease (Terpios hoshinota): A Probable Cause for the Rapid Coral Mortality at the Northern Reef of Yongxing Island in the South China Sea. Ambio 41:446–455.

12. Madduppa H, Schupp PJ, Faisal MR, Sastria MY, Thoms C. 2017. Persistent outbreaks of the "black disease" sponge Terpios hoshinota in Indonesian coral reefs. Marine Biodiversity 47:149–151.

13. Hentschel U, Piel J, Degnan SM, Taylor MW. 2012. Genomic insights into the marine sponge microbiome. Nature Reviews Microbiology 10:641–U75.

14. Tang SL, Hong MJ, Liao MH, Jane WN, Chiang PW, Chen CB, Chen CA. 2011. Bacteria associated with an encrusting sponge (Terpios hoshinota) and the corals partially covered by the sponge. Environmental Microbiology 13:1179–1191.

15. Hirose E, Murakami A. 2011. Microscopic anatomy and pigment characterization of coral-encrusting black sponge with cyanobacterial symbiont, Terpios hoshinota. Zoolog Sci 28:199–205.

16. Wang JT, Hirose E, Hsu CM, Chen YY, Meng PJ, Chen CA. 2012. A Coral-Killing Sponge, Terpios hoshinota, Releases Larvae Harboring Cyanobacterial Symbionts: An Implication of Dispersal. Zoological Studies 51:314–320.

17. Hsu CM, Wang JT, Chen CA. 2013. Larval release and rapid settlement of the coral- killing sponge, Terpios hoshinota, at Green Island, Taiwan. Marine Biodiversity 43:259–260.

18. Nozawa Y, Huang YS, Hirose E. 2016. Seasonality and lunar periodicity in the sexual reproduction of the coral-killing sponge, Terpios hoshinota. Coral Reefs 35:1071–1081.

19. Soong K, Yang SL, Chen CA. 2009. A Novel Dispersal Mechanism of a Coral- Threatening Sponge, Terpios hoshinota (Suberitidae, Porifera). Zoological Studies 48:596–596.

20. Thinesh T, Meenatchi R, Pasiyappazham R, Jose PA, Selvan M, Kiran GS, Selvin J. 2017. Short-term in situ shading effectively mitigates linear progression of coral- killing sponge Terpios hoshinota. Plos One 12.

21. Yu CH, Lu CK, Su HM, Chiang TY, Hwang CC, Liu T, Chen YM. 2015. Draft genome of Myxosarcina sp. strain GI1, a baeocytous cyanobacterium associated with the marine sponge Terpios hoshinota. Stand Genomic Sci 10:28.

22. Webster NS, Thomas T. 2016. The Sponge Hologenome. mBio 7:e00135–16.

23. Pita L, Rix L, Slaby BM, Franke A, Hentschel U. 2018. The sponge holobiont in a changing ocean: from microbes to ecosystems. Microbiome 6:46.

24. Pita L, Turon X, Lopez-Legentil S, Erwin PM. 2013. Host rules: spatial stability of bacterial communities associated with marine sponges (Ircinia spp.) in the Western Mediterranean Sea. FEMS Microbiol Ecol 86:268–76.

25. Whatley JM. 1977. The fine structure of Prochloron. New Phytologist 79:309–313.

26. Kolmogorov M, Bickhart DM, Behsaz B, Gurevich A, Rayko M, Shin SB, Kuhn K, Yuan J, Polevikov E, Smith TPL, Pevzner PA. 2020. metaFlye: scalable long-read metagenome assembly using repeat graphs. Nat Methods 17:1103–1110.

27. Podell S, Blanton JM, Oliver A, Schorn MA, Agarwal V, Biggs JS, Moore BS, Allen EE. 2020. A genomic view of trophic and metabolic diversity in clade-specific Lamellodysidea sponge microbiomes. Microbiome 8:97.

28. Richter M, Rossello-Mora R. 2009. Shifting the genomic gold standard for the prokaryotic species definition. Proc Natl Acad Sci U S A 106:19126–31.

29. Qin QL, Xie BB, Zhang XY, Chen XL, Zhou BC, Zhou J, Oren A, Zhang YZ. 2014. A proposed genus boundary for the prokaryotes based on genomic insights. J Bacteriol 196:2210–5.

30. Kuhl M, Behrendt L, Trampe E, Qvortrup K, Schreiber U, Borisov SM, Klimant I, Larkum AW. 2012. Microenvironmental Ecology of the Chlorophyll b-Containing Symbiotic Cyanobacterium Prochloron in the Didemnid Ascidian Lissoclinum patella. Front Microbiol 3:402.

31. Harding K, Turk-Kubo KA, Sipler RE, Mills MM, Bronk DA, Zehr JP. 2018. Symbiotic unicellular cyanobacteria fix nitrogen in the Arctic Ocean. Proc Natl Acad Sci U S A 115:13371–13375.

32. Mehbub MF, Perkins MV, Zhang W, Franco CMM. 2016. New marine natural products from sponges (Porifera) of the order Dictyoceratida (2001 to 2012); a promising source for drug discovery, exploration and future prospects. Biotechnol Adv 34:473–491.

33. Schorn MA, Jordan PA, Podell S, Blanton JM, Agarwal V, Biggs JS, Allen EE, Moore BS. 2019. Comparative Genomics of Cyanobacterial Symbionts Reveals Distinct, Specialized Metabolism in Tropical Dysideidae Sponges. mBio 10.

34. Katayama T, Ozaki S, Keyamura K, Fujimitsu K. 2010. Regulation of the replication cycle: conserved and diverse regulatory systems for DnaA and oriC. Nat Rev Microbiol 8:163–70.

35. Ohbayashi R, Hirooka S, Onuma R, Kanesaki Y, Hirose Y, Kobayashi Y, Fujiwara T, Furusawa C, Miyagishima SY. 2020. Evolutionary Changes in DnaA-Dependent Chromosomal Replication in Cyanobacteria. Front Microbiol 11:786.

36. Ohbayashi R, Watanabe S, Ehira S, Kanesaki Y, Chibazakura T, Yoshikawa H. 2016. Diversification of DnaA dependency for DNA replication in cyanobacterial evolution. ISME J 10:1113–21.

37. Lesser MP, Fiore C, Slattery M, Zaneveld J. 2016. Climate change stressors destabilize the microbiome of the Caribbean barrel sponge, Xestospongia muta. Journal of Experimental Marine Biology and Ecology 475:11–18.

38. Morrow KM, Fiore CL, Lesser MP. 2016. Environmental drivers of microbial community shifts in the giant barrel sponge, Xestospongia muta, over a shallow to mesophotic depth gradient. Environ Microbiol 18:2025–38.

39. Webster NS, Reusch TBH. 2017. Microbial contributions to the persistence of coral reefs. ISME J 11:2167–2174.

40. Nguyen MT, Liu M, Thomas T. 2014. Ankyrin-repeat proteins from sponge symbionts modulate amoebal phagocytosis. Mol Ecol 23:1635–45.

41. Wiens M, Korzhev M, Krasko A, Thakur NL, Perovic-Ottstadt S, Breter HJ, Ushijima H, Diehl-Seifert R, Muller IM, Muller WEG. 2005. Innate immune defense of the sponge Suberites domuncula against bacteria involves a MyD88-dependent signaling pathway - Induction of a perforin-like molecule. Journal of Biological Chemistry 280:27949–27959.

42. Fan L, Reynolds D, Liu M, Stark M, Kjelleberg S, Webster NS, Thomas T. 2012. Functional equivalence and evolutionary convergence in complex communities of microbial sponge symbionts. Proceedings of the National Academy of Sciences of the United States of America 109:E1878–E1887.

43. Reveillaud J, Maignien L, Eren AM, Huber JA, Apprill A, Sogin ML, Vanreusel A. 2014. Host-specificity among abundant and rare taxa in the sponge microbiome. Isme Journal 8:1198–1209.

44. Burgsdorf I, Erwin PM, Lopez-Legentil S, Cerrano C, Haber M, Frenk S, Steindler L. 2014. Biogeography rather than association with cyanobacteria structures symbiotic microbial communities in the marine sponge Petrosia ficiformis. Frontiers in Microbiology 5.

45. Gao ZM, Wang Y, Tian RM, Wong YH, Batang ZB, Al-Suwailem AM, Bajic VB, Qian PY. 2014. Symbiotic Adaptation Drives Genome Streamlining of the Cyanobacterial Sponge Symbiont "Candidatus Synechococcus spongiarum". Mbio 5.

46. Lo WS, Huang YY, Kuo CH. 2016. Winding paths to simplicity: genome evolution in facultative insect symbionts. Fems Microbiology Reviews 40:855–874.

47. Moran NA, McCutcheon JP, Nakabachi A. 2008. Genomics and Evolution of Heritable Bacterial Symbionts. Annual Review of Genetics 42:165–190.

48. Kolman MA, Nishi CN, Perez-Cenci M, Salerno GL. 2015. Sucrose in cyanobacteria: from a salt-response molecule to play a key role in nitrogen fixation. Life (Basel) 5:102–26.

49. Curatti L, Giarrocco LE, Cumino AC, Salerno GL. 2008. Sucrose synthase is involved in the conversion of sucrose to polysaccharides in filamentous nitrogen-fixing cyanobacteria. Planta 228:617–625.

50. Blumwald E, Telor E. 1982. Osmoregulation and Cell Composition in Salt- Adaptation of Nostoc-Muscorum. Archives of Microbiology 132:168–172.

51. Desplats P, Folco E, Salerno GL. 2005. Sucrose may play an additional role to that of an osmolyte in Synechocystis sp PCC 6803 salt-shocked cells. Plant Physiology and Biochemistry 43:133–138.

52. Ran L, Larsson J, Vigil-Stenman T, Nylander JA, Ininbergs K, Zheng WW, Lapidus A, Lowry S, Haselkorn R, Bergman B. 2010. Genome erosion in a nitrogen-fixing vertically transmitted endosymbiotic multicellular cyanobacterium. PLoS One 5:e11486.

53. Akman L, Yamashita A, Watanabe H, Oshima K, Shiba T, Hattori M, Aksoy S. 2002. Genome sequence of the endocellular obligate symbiont of tsetse flies, Wigglesworthia glossinidia. Nat Genet 32:402–7.

54. Frank AC. 2019. Molecular host mimicry and manipulation in bacterial symbionts. FEMS Microbiol Lett 366.

55. Taylor MW, Radax R, Steger D, Wagner M. 2007. Sponge-associated microorganisms: evolution, ecology, and biotechnological potential. Microbiol Mol Biol Rev 71:295–347.

56. Wilkinson CR. 1983. Net primary productivity in coral reef sponges. Science 219:410–2.

57. Wang JT, Hsu CM, Kuo CY, Meng PJ, Kao SJ, Chen CA. 2015. Physiological Outperformance at the Morphologically-Transformed Edge of the Cyanobacteriosponge Terpios hoshinota (Suberitidae: Hadromerida) when Confronting Opponent Corals. Plos One 10.

58. Zhang F, Blasiak LC, Karolin JO, Powell RJ, Geddes CD, Hill RT. 2015. Phosphorus sequestration in the form of polyphosphate by microbial symbionts in marine sponges. Proceedings of the National Academy of Sciences of the United States of America 112:4381–4386.

59. Colman AS. 2015. Sponge symbionts and the marine P cycle. Proceedings of the National Academy of Sciences of the United States of America 112:4191–4192.

60. Utami RT, Zamani NP, Madduppa HH. 2018. Molecular identification, abundance and distribution of the coral-killing sponge Terpios hoshinota in Bengkulu and Seribu Islands, Indonesia. Biodiversitas Journal of Biological Diversity 19:2238–2246.

61. Bryan PG. 1973. Growth rate, toxicity and distribution of the encrusting sponge Terpios sp.(Hadromerida: Suberitidae) in Guam, Mariana Islands. Micronesica 9:237–242.

62. Thinesh T, Meenatchi R, Lipton AN, Anandham R, Jose PA, Tang SL, Seghal Kiran G, Selvin J. 2020. Metagenomic sequencing reveals altered bacterial abundance during coral-sponge interaction: Insights into the invasive process of coral-killing sponge Terpios hoshinota. Microbiol Res 240:126553.

63. Teruya T, Nakagawa S, Koyama T, Arimoto H, Kita M, Uemura D. 2004. Nakiterpiosin and nakiterpiosinone, novel cytotoxic C-nor-D-homosteroids from the Okinawan sponge Terpios hoshinota. Tetrahedron 60:6989–6993.

64. Teruya T, Nakagawa S, Koyama T, Suenaga K, Uemura D. 2002. Terpiodiene: A novel tricyclic alcohol from the Okinawan sponge Terpios hoshinota. Chemistry Letters doi:DOI 10.1246/cl.2002.38:38-39.

65. Fagundes MB, Falk RB, Facchi MMX, Vendruscolo RG, Maroneze MM, Zepka LQ, Jacob-Lopes E, Wagner R. 2019. Insights in cyanobacteria lipidomics: A sterols characterization from Phormidium autumnale biomass in heterotrophic cultivation. Food Research International 119:777–784.

66. Lane D. 1991. 16S/23S rRNA sequencing. Nucleic acid techniques in bacterial systematics:115–175.

67. Schloss PD, Westcott SL, Ryabin T, Hall JR, Hartmann M, Hollister EB, Lesniewski RA, Oakley BB, Parks DH, Robinson CJ, Sahl JW, Stres B, Thallinger GG, Van Horn DJ, Weber CF. 2009. Introducing mothur: Open-Source, Platform-Independent, Community-Supported Software for Describing and Comparing Microbial Communities. Applied and Environmental Microbiology 75:7537–7541.

68. Edgar RC. 2010. Search and clustering orders of magnitude faster than BLAST. Bioinformatics 26:2460–2461.

69. Edgar RC. 2013. UPARSE: highly accurate OTU sequences from microbial amplicon reads. Nature Methods 10:996-+.

70. Yilmaz P, Parfrey LW, Yarza P, Gerken J, Pruesse E, Quast C, Schweer T, Peplies J, Ludwig W, Glockner FO. 2014. The SILVA and "All-species Living Tree Project (LTP)" taxonomic frameworks. Nucleic Acids Research 42:D643–D648.

71. Bolyen E, Rideout JR, Dillon MR, Bokulich N, Abnet CC, Al-Ghalith GA, Alexander H, Alm EJ, Arumugam M, Asnicar F, Bai Y, Bisanz JE, Bittinger K, Brejnrod A, Brislawn CJ, Brown CT, Callahan BJ, Caraballo-Rodriguez AM, Chase J, Cope EK, Da Silva R, Diener C, Dorrestein PC, Douglas GM, Durall DM, Duvallet C, Edwardson CF, Ernst M, Estaki M, Fouquier J, Gauglitz JM, Gibbons SM, Gibson DL, Gonzalez A, Gorlick K, Guo JR, Hillmann B, Holmes S, Holste H, Huttenhower C, Huttley GA, Janssen S, Jarmusch AK, Jiang LJ, Kaehler BD, Bin Kang K, Keefe CR, Keim P, Kelley ST, Knights D, et al. 2019. Reproducible, interactive, scalable and extensible microbiome data science using QIIME 2. Nature Biotechnology 37:852–857.

72. Chaumeil PA, Mussig AJ, Hugenholtz P, Parks DH. 2020. GTDB-Tk: a toolkit to classify genomes with the Genome Taxonomy Database. Bioinformatics 36:1925–1927.

73. Bolger AM, Lohse M, Usadel B. 2014. Trimmomatic: a flexible trimmer for Illumina sequence data. Bioinformatics 30:2114–2120.

74. Walker BJ, Abeel T, Shea T, Priest M, Abouelliel A, Sakthikumar S, Cuomo CA, Zeng Q, Wortman J, Young SK, Earl AM. 2014. Pilon: an integrated tool for comprehensive microbial variant detection and genome assembly improvement. PLoS One 9:e112963.

75. Seemann T. 2014. Prokka: rapid prokaryotic genome annotation. Bioinformatics 30:2068–9.

76. Tatusov RL, Galperin MY, Natale DA, Koonin EV. 2000. The COG database: a tool for genome-scale analysis of protein functions and evolution. Nucleic Acids Research 28:33–36.

77. Kanehisa M, Goto S. 2000. KEGG: Kyoto Encyclopedia of Genes and Genomes.Nucleic Acids Research 28:27–30.

78. Kanehisa M, Sato Y, Morishima K. 2016. BlastKOALA and GhostKOALA: KEGG Tools for Functional Characterization of Genome and Metagenome Sequences. Journal of Molecular Biology 428:726–731.

79. Hyatt D, Chen GL, LoCascio PF, Land ML, Larimer FW, Hauser LJ. 2010. Prodigal: prokaryotic gene recognition and translation initiation site identification. Bmc Bioinformatics 11.

80. Joel A Boyd, BJW, GWT. 2019. Comparative genomics using EnrichM. In preparation.

81. Kanehisa M, Sato Y. 2020. KEGG Mapper for inferring cellular functions from protein sequences. Protein Science 29:28–35.

82. Camacho C, Coulouris G, Avagyan V, Ma N, Papadopoulos J, Bealer K, Madden TL. 2009. BLAST plus : architecture and applications. Bmc Bioinformatics 10.

83. Elbourne LDH, Tetu SG, Hassan KA, Paulsen IT. 2017. TransportDB 2.0: a database for exploring membrane transporters in sequenced genomes from all domains of life. Nucleic Acids Research 45:D320–D324.

84. Coordinators NR. 2018. Database resources of the National Center for Biotechnology Information. Nucleic Acids Res 46:D8–D13.

85. Edgar RC. 2004. MUSCLE: a multiple sequence alignment method with reduced time and space complexity. BMC Bioinformatics 5:113.

86. Kumar S, Stecher G, Tamura K. 2016. MEGA7: Molecular Evolutionary Genetics Analysis Version 7.0 for Bigger Datasets. Mol Biol Evol 33:1870–4.

87. Minh BQ, Schmidt HA, Chernomor O, Schrempf D, Woodhams MD, von Haeseler A, Lanfear R. 2020. IQ-TREE 2: New Models and Efficient Methods for Phylogenetic Inference in the Genomic Era. Mol Biol Evol 37:1530–1534.

88. Letunic I, Bork P. 2019. Interactive Tree Of Life (iTOL) v4: recent updates and new developments. Nucleic Acids Res 47:W256–W259.

89. Parks DH, Chuvochina M, Waite DW, Rinke C, Skarshewski A, Chaumeil PA, Hugenholtz P. 2018. A standardized bacterial taxonomy based on genome phylogeny substantially revises the tree of life. Nature Biotechnology 36:996-+.

